# Impact of sustained TGFβ receptor inhibition on chromatin accessibility and gene expression in cultured human endometrial MSC

**DOI:** 10.1101/2020.05.01.073346

**Authors:** Raffaella Lucciola, Pavle Vrljicak, Caitlin Filby, Saeedeh Darzi, Shanti Gurung, Joanne Muter, Sascha Ott, Jan J Brosens, Caroline E Gargett

## Abstract

Endometrial mesenchymal stem cells (eMSC) drive the extraordinary regenerative capacity of the human endometrium. Clinical application of eMSC for therapeutic purposes is hampered by spontaneous differentiation and cellular senescence upon large-scale expansion *in vitro*. A83-01, a selective transforming growth factor-β receptor (TGFβ-R) inhibitor, promotes pharmacological expansion of eMSC in culture by blocking differentiation and senescence, but the underlying mechanisms are incompletely understood. In this study, we combined RNA-seq and ATAC-seq to study the impact of sustained TGFβ-R inhibition on gene expression and chromatin architecture of eMSC. Treatment of primary eMSC with A83-01 for 5 weeks resulted in differential expression of 1,463 genes. Gene ontology analysis showed enrichment of genes implicated in cell growth whereas extracellular matrix genes and genes involved in cell fate commitment were downregulated. ATAC-seq analysis demonstrated that sustained TGFβ-R inhibition results in opening and closure of 3,555 and 2,412 chromatin loci, respectively. Motif analysis revealed marked enrichment of retinoic acid receptor (RAR) binding sites, which was paralleled by the induction of *RARB*, encoding retinoic acid receptor beta (RARβ). Selective RARβ inhibition attenuated proliferation and clonogenicity of A83-01 treated eMSC. Taken together, our study provides new insights into the gene networks and genome-wide chromatin changes that underpin maintenance of an undifferentiated phenotype of eMSC in prolonged culture.

**Significance statement:** Cycling human endometrium is a rich source of adult stem/progenitor cells that could be exploited for clinical purposes. Small molecules, such as A83-01, that modulate cell identity may open new avenues to maintain the functional properties of eMSC upon expansion in culture. By integrating complementary genome-wide profiling techniques, we mapped the dynamic changes in chromatin landscape and gene expression in response to prolonged A83-01 treatment of eMSC. Our findings provide new insights into the mechanisms of action of TGFβ-R inhibition that may lead to the development of more targeted pharmacological approaches for MSC expansion.

## Introduction

The human endometrium is a highly dynamic tissue that generates 4-10 mm of mucosa in each menstrual cycle^1^. This extraordinary regenerative capacity is mediated by resident epithelial progenitors and mesenchymal stem/stromal cells (MSC)^2, 3^. Cultured endometrial MSC (eMSC) are clonogenic, highly proliferative, multipotent, and express the International Society for Cellular Therapies (ISCT) surface markers.^4^ Human eMSC can be purified as CD140b^+^CD146^+^ cells^5^, or by means of the surface marker SUSD2 (Sushi Domain-containing 2, formerly W5C5).^6^ While eMSC also express Stro-1, this marker does not enrich for clonogenic cells.^7^ eMSC further express MSCA-1^8^ and foremost reside in the perivasculature of spiral arterioles and venules.^9–11^ A major advantage of the endometrium over other sources of adult MSC, such as bone marrow or adipose tissue, is its accessibility. Endometrial sampling is a routine office-based procedure that does not require anesthesia.^9^

Preclinical animal studies showed that eMSC are a promising cell source for treatment of gynecological disorders, including pelvic organ prolapse.^12, 13^ For example, eMSC seeded or bio-printed onto meshes with biomechanical properties matching the human vagina (e.g. non-degradable polyamide/gelatin composite meshes),^14, 15 16^, or on degradable nanofibers,^17^ promote angiogenesis, collagen deposition, and cellular infiltration into biomaterials when transplanted into rodent or ovine models. They also elicit an early inflammatory response, characterized first by influx of M1 macrophages, which then switch to the M2 wound healing phenotype.^18, 19^ *In vitro,* eMSC seeded on polyamide/gelatin meshes differentiate into smooth muscle cells and fibroblasts; cell types that are important for restoring vaginal structure and function.^20^ Hence, the endometrium is a rich source of MSC for autologous and allogenic cell-based therapies, including pelvic organ prolapse, urinary incontinence, and regeneration of scarred endometrium in women with Asherman’s syndrome.^2, 21, 22^

Clinical application of eMSC requires expansion of cells in culture.^12, 23^ As is the case for MSC from other sources,^24^ eMSC cultured over several passages differentiate spontaneously and are subjected to replicative stress caused by telomere shortening.^23, 25, 26^ Consequently, the cells lose their proliferative capacity as well as the ability to reconstitute tissue *in vivo.*^24^ To overcome this impediment to the clinical translation, we and others have explored the use of small molecules to maintain the *in vivo* properties of eMSC and MSC in prolonged culture.^23, 25, 27^ We recently reported that A83-01, a selective inhibitor of TGF-β type I receptor (TGFβ-R) ALK4, 5 and 7 kinase, increases proliferation and inhibits apoptosis and senescence of cultured eMSC, thereby safeguarding their functional properties *in vitro*^23^ and prolonging their survival *in vivo.*^28^ Further, A83-01-treated eMSC exhibit increased angiogenic activity and express a proangiogenic, antifibrotic, and immunomodulatory gene profile.^23, 25^

The mechanisms underlying pharmacological expansion of cultured eMSC in response to sustained TGFβ-R inhibition are incompletely understood. Adult stem/progenitor cells have more open chromatin than their differentiated progeny, but less than pluripotent stem cells. Further dynamic chromatin changes underpin subsequent differentiation into mesodermal lineages.^29^ Based on these observations, we hypothesized that sustained TGFβ-R inhibition leads to divergence in the chromatin landscape of cultured eMSC and activation of transcriptional regulatory circuitries that maintain the cells in a more naïve or undifferentiated state. To test this hypothesis, eMSC cultured with or without A83-01 were subjected to integrated RNA-sequencing (RNA-seq) and Assay for Transposase Accessible Chromatin through sequencing (ATAC-seq).

## Materials and Methods

### Endometrial biopsies

The study was approved by the Monash Health and Monash University Human Research Ethics committees. Endometrial biopsies were obtained from seven pre-menopausal women, without endometrial pathologies, following written informed consent and according to The Declaration of Helsinki (2000) guidelines. Participant information is kept confidential and samples deidentified prior to use. None of the participants received hormonal treatment within three months prior to the biopsy.

### eMSC isolation and culture

Endometrial biopsies were processed and single-cell suspensions of eMSC obtained as described previously.^6^ Briefly, finely minced endometrial tissue was enzymatically and mechanically digested in Dulbecco’s modified Eagle’s medium (DMEM/F12) supplemented with collagenase type I and DNase I (Worthington Biochemical Corporation, USA) at 37°C on a rotating MACSmix (Miltenyi Biotec, USA) for 60 min. The digested tissue was filtered through 40 μm strainers (Becton Dickinson, BD, USA) to remove gland fragments, washed, red blood cells removed using Ficoll-Paque (GE healthcare Bio-science, USA) density gradient separation at 1500 rpm for 15 min, and the interface containing stromal cells was washed in DMEM/F12/ 10% fetal calf serum (FCS, Invitrogen, USA) and 1% primocin (Life Technologies, USA). Single-cell suspensions were re-suspended in separation buffer (0.5% FCS/PBS), and incubated with 10 μg/ml phycoerythrin (PE)-conjugated anti-human SUSD2 (BioLegend, USA) in the dark at 4°C for 30 min. Cells were then washed, the cell pellet re-suspended in 20 μl of anti-PE magnetic-activated cell sorting (MACS) microbeads (Miltenyi Biotec) with 80 μL separation buffer and incubated in the dark at 4°C for 30 min. The cells were washed, re-suspended in separation buffer 500 μL and applied to a Miltenyi column (Miltenyi Biotech) in a magnetic field. Columns were washed three times with buffer. Magnetically labelled SUSD2^+^ eMSC were eluted with buffer and cell number was determined using Koya glasstic slides (KOVA International, USA).

eMSC were cultured in DMEM/F12 medium containing 10% FCS, 1% primocin and 2mM glutamine (Invitrogen), supplemented with 10 ng/ml basic fibroblast growth factor (bFGF) (Peprotech, USA) and scaled-down to an in-house DMEM/F12 serum free medium (SFM) over 48 h within the first passage and incubated at 37°C in 5% CO_2_/5% O_2_/90% N_2_, as described previously ^30^, with 1 μM TGFβ-R inhibitor, A83-01 (Tocris Bioscience, USA) or vehicle control (0.01% DMSO). Cells were seeded at 5,000 cells/cm^2^ with medium changed every 48 h and passaged on days 15, 22, 29 and 36 into fibronectin-coated (10 μg/ml; BD, USA) culture flasks as described previously ^23^. At each passage, cells were counted using Kova glasstic slides and cumulative cell population (total cell number) calculated by multiplying total number of cells yielded at the current passage by total number of cells yielded at the previous passage and then dividing by the number of cells seeded at the current passage, as described previously.^31^ At passage 4 (i.e. 36 days in culture), untreated and A83-01-treated eMSC cultures were subjected to functional assays, and RNA- and ATAC-seq.

### Flow cytometry

The surface phenotype of untreated and A83-01-treated eMSC was assessed by flow cytometry for three MSC markers [SUSD2, CD140b (Platelet-derived growth factor receptor β) and CD90 (Thy-1)] as described previously.^23^ Cells were incubated for 1 h with 1:20 primary or matched isotype control antibodies in PBS containing 2% FCS at 4°C in dark. The primary antibodies were APC-conjugated SUSD2 (Biolegend), PE-conjugated CD140b (R&D Systems, USA) and APC-conjugated CD90 (BD Pharmingen, USA). Cells were washed with PBS/2% FCS and fixed with 4% paraformaldehyde in PBS/2% FCS. eMSC were analysed using BD FACSCanto II (BD) (10,000 events /sample) and FlowJo v.10 software.

### Colony-forming unit-fibroblast (CFU-F) assay

eMSC were cultured with or without A83-01 for 2 passages (29 days) and then seeded at 50 cells/cm^2^ to assess cloning efficiency as described previously,^23^ with minor modifications. Briefly, eMSC were seeded on fibronectin-coated 10 mm culture dishes and cultured in SFM, supplemented with bFGF and EGF (both 10 ng/mL; Invitrogen), in the presence or absence of A83-01 (1 μM) in SFM at 37°C in 5% CO_2_5% O_2_/90% N_2_ for 2 weeks, with weekly medium changes. Cultures were then fixed in 10% formalin and stained with hematoxylin (Amber Scientific, USA), washed and blued in Scott’s tap water. Colonies were counted and colony efficiency determined by dividing total number of colonies by number of cells seeded and multiplied by 100.

### RNA extraction, RNA quality control and RNA libraries

Total RNA was extracted from untreated and A83-01-treated eMSC using RNeasy Mini kit (Qiagen, Germany) according to the manufacturer’s instructions, with some variations. Briefly, eMSC were trypsinised using TryLE™ (Life Technologies), resuspended in DMEM/F12 containing 5% Albumax and then centrifuged at 300 g for 5 min. Cell were lysed with RNeasy Lysis Buffer, genomic DNA contamination removed with RNase-free DNase (Qiagen) and RNA eluted with 30-50 μl RNase-free water. RNA quality was assessed on an Agilent Technologies 2100 Bioanalyzer according to the manufacturer’s instructions. Samples with RNA Integrity Number (RIN) >8 were subjected TruSeq Poly-A mRNA Library Pro Kit protocol 15031047 RevD (Illumina, USA) to generate indexed cDNA libraries. The library size was assessed on an Agilent Bioanalyzer and quantified by Qubit and qPCR.

### RNA-seq

RNA libraries were sequenced by Illumina HiSeq3000. Fifty million single-end reads were sequenced per sample with a read length of 50 bp. Transcriptomic maps were identified using Bowtie-2.2.3 ^32^ and Samtools-1.2.0^33^ against the UCSC hg19 transcriptome reference from the Illumina iGenomes resource (2014). Counts were assessed using HTSeq-0.6.1^34^ and transcripts per million (TPM) were calculated. DESeq2^35^ was used for detection of differentially expressed genes in a pair-wise manner. Differentially expressed genes were subjected to Gene Ontology (GO) and KEGG Pathway enrichment analyses using the Database for Annotation, Visualization and Integrated Discovery (DAVID) version 6.8^36^ and visualized using Reduce and Visualize Gene Ontology (REVIGO) online software.^37^ Fastq, metadata spreadsheet and table of counts have been deposited in the National Centre for Biotechnology Information Gene Expression Omnibus/sequence Read Archive with GEO accession number GSE146067

### ATAC-seq

ATAC-seq was performed as described^38^, although with some modifications.^39^ Briefly, cells were washed with cold PBS, lysed using cold EZ lysis buffer (10 mM Tris-HCl, pH 7.4, 10 mM NaCl, 3 mM MgCl_2_ and 0.1% IGEPAL CA-630, Sigma-Aldrich, UK), transferred to chilled nuclease-free tubes, vortexed, left on ice for 5 minutes, and then pelleted in a refrigerated centrifuge. The nuclear pellet was washed in EZ lysis buffer and re-suspended in the transposase reaction mix containing 25 μl Tagment DNA (TD) Buffer, 5 μl Tagment DNA Enzyme and 20 μl nuclease free water (Nextera DNA Sample Preparation Kit, Illumina, UK) for 45 min at 37°C. Samples were purified using a Zymo DNA Clean and Concentrator-5 Purification kit (Zymo Research, USA). Briefly, DNA binding buffer was added to 50 μl samples, mixed and transferred to the column, centrifuged at 17,000*g* for 30 sec at RT, 200 μl DNA wash buffer added, columns centrifuged and repeated twice. After removing the residual liquid, 23 μl pre-warmed elution buffer was added and incubated for 2 min at RT, then centrifuged for 2 min to elute DNA. 20 μl samples were added to PCR tubes containing 5 μl index 1, 5 μl index 2, 15 μl Master mix (NPM), 5 μl Primer Cocktail (Nextera DNA Sample Preparation Kit and Nextera Index Kit, Illumina). Amplification was performed in a Veriti 96 Well Thermal Cycler (Applied Biosystems, USA) using the following PCR conditions: 72°C for 3 min, 98°C for 30 sec then 15 cycles of 98°C for 10 sec, 63°C for 30 sec, 72°C for 1 min. Libraries were purified using AMPure XP beads using the Illumina Nextera kit recommended protocol and quantified using Qubit HS DNA Assay on a Qubit 2.0 Fluorometer. Library sizes were assessed by Agilent Bioanalyzer using the High Sensitivity DNA chip. ATAC-seq library samples were sequenced on an Illumina HiSeq 1500 to a depth of thirty million paired-end reads/sample, with a read length of 100 bp.

### ATAC-seq data and motif analyses

Sequenced paired-end reads were aligned to the University of California Santa Cruz (UCSC) human genome 19 (hg19) assembly using Bowtie2-2.2.6^32^ and Samtools-1.2.0^33^ and peak calling performed using MACS-2.1.0. HTSeq-0.6.1^34^ to count the reads overlapping the peaks and differential expression analysis of sequencing data 2 (DESeq2) was used to determine opening and closing regions of the chromatin.^35^ Fastq, metadata spreadsheet and table of counts have been deposited in the National Centre for Biotechnology Information Gene Expression Omnibus/sequence Read Archive with GEO accession number GSE146067. Differential open chromatin regions were mapped to *cis*-regulatory elements of their proximal genes using ENCODE DNaseI hypersensitivity data.^40^ Physical interaction and distance no greater than 10 kb were used as criteria to assess association between ATAC-seq peak and proximal gene regulatory element. *De novo* short sequence motif analysis using Hypergeometric Optimization of Motif EnRichment (HOMER) v.4.8 was performed on 3,555 opening and 2,412 closing ATAC-seq peaks to determine enrichment and depletion of TF short sequence binding motifs in the differential ATAC-seq peaks.^41^

### RARβ inhibition experiments

Passage 3 cultured eMSC from 4 biological samples were treated with 1 μM A83-01 or 0.01% DMSO (vehicle control) for 7 days, then trypsinized and seeded in triplicate at 1000 cells/well (3.125×10^3^ cells/cm^2^) into fibronectin-coated wells of two 96 well plates per biological replicate to assess proliferation on day 0 (D0) and day 3 (D3) in the presence or absence of 10 μM RARβ antagonist LE135 (Tocris Bioscience) using the cell viability MTS assay. After seeding, cells were allowed to adhere for 1 h before addition of 100 μl medium containing A83-01+LE135 or A83-01+DMSO. After 2 h, the MTS reagent (20μl, CellTiter 96® AQueous One Solution Cell Proliferation Assay, Promega, USA) was added to wells for the D0 timepoint and incubated for 2.25 h in the dark, then absorbance read at 490 nm on a spectrophotometer (Spectramax i3, Molecular Devices, USA). The medium was changed on the D3 plate at 24 h and the viability assay completed 72 h after seeding, as described above. Parallel cultures were subjected to CFU-F assays. Data were corrected for background readings (medium only) for each plate and normalized to D0 A83-01 for each sample and reported as fold-change.

### Statistical Analysis

Statistical analyses were performed with GraphPad Prism 8. Technical replicates were inspected for outliers, which were removed from the analysis using Grubbs test. Normality of the data was determined with Shapiro-Wilk normality test. Individual data points and mean ± standard error of the mean (SEM) are shown when appropriate. Statistical significance was determined using two-way analysis of variance (ANOVA) or ratio paired *t*-tests with *p*<0.05 considered statistically significant. For RNA-seq data analysis, statistical significance was assessed using the Benjamini-Hochberg procedure to control false discovery rate. Changes in gene expression were deemed statistically significant if the adjusted *p*-value (*q*-value) was less than 0.05.

## Results

### Phenotypic characterization of sustained A83-01-treated eMSC

To investigate if eMSC can be expanded more efficiently when maintained under sustained TGFβ-R inhibition from culture initiation, cells isolated from 3 individual biopsies were seeded at 5000 cells/cm^2^ in SFM supplemented with either A83-01 or vehicle (DMSO). The medium was refreshed every 48 h and cumulative cell population calculated at each passage (culture days 15, 22, 29 and 36). As shown in Figure 1A, treatment with A83-01 from culture initiation progressively conferred a proliferation advantage. After 36 days in culture, at least one order of magnitude more cells were produced in the A83-01 medium when compared to control medium (Fig. 1A). Next, we performed colony-forming unit-fibroblast (CFU-F) assays on eMSC first cultured with or without A83-01 for 2 passages (29 days) and then seeded at a low density (50 cells/cm2) to allow colony formation for a further 14 days. In keeping with our previous study,^23^ exposure of cultured eMSC to A83-01 increased the CFU-F activity of primary cultures between ~ 4-10 fold (*p* = 0.0146; Fig. 1B). We then chose the 36-day timepoint (Fig. 1A) to analyze A83-01-treated cells for the expression of phenotypic eMSC markers CD140b, SUSD2 and CD90 by flow cytometry. The abundance of CD90^+^ cells was 99.1±0.5% (n=3) for the control and did not change upon TGFβ-R blockade (99.7±0.2% (n=3) (Fig. 1C). By contrast, A83-01 treatment consistently increased the abundance of CD140b^+^ cells (*p*=0.031) and SUSD2^+^ cells (*p*=0.078) (Fig. 1C), although not statistically significant for SUSD2. Mean fluorescence intensity (MFI) was also calculated to evaluate the abundance of different cell surface molecules/cell. The MFI for CD90 did not show a consistent change upon A83-01 treatment, in keeping with previous observations that this ISCT minimal criteria MSC marker^42^ shows little change under varied culture conditions.^23, 30^ By contrast, the MFI of CD140b and SUSD2 (Fig. 1D) increased in response to A83-01 treatment in each culture, although statistical significance was not reached (*p*=0.0715 and 0.0897, respectively).

**Figure 1.**
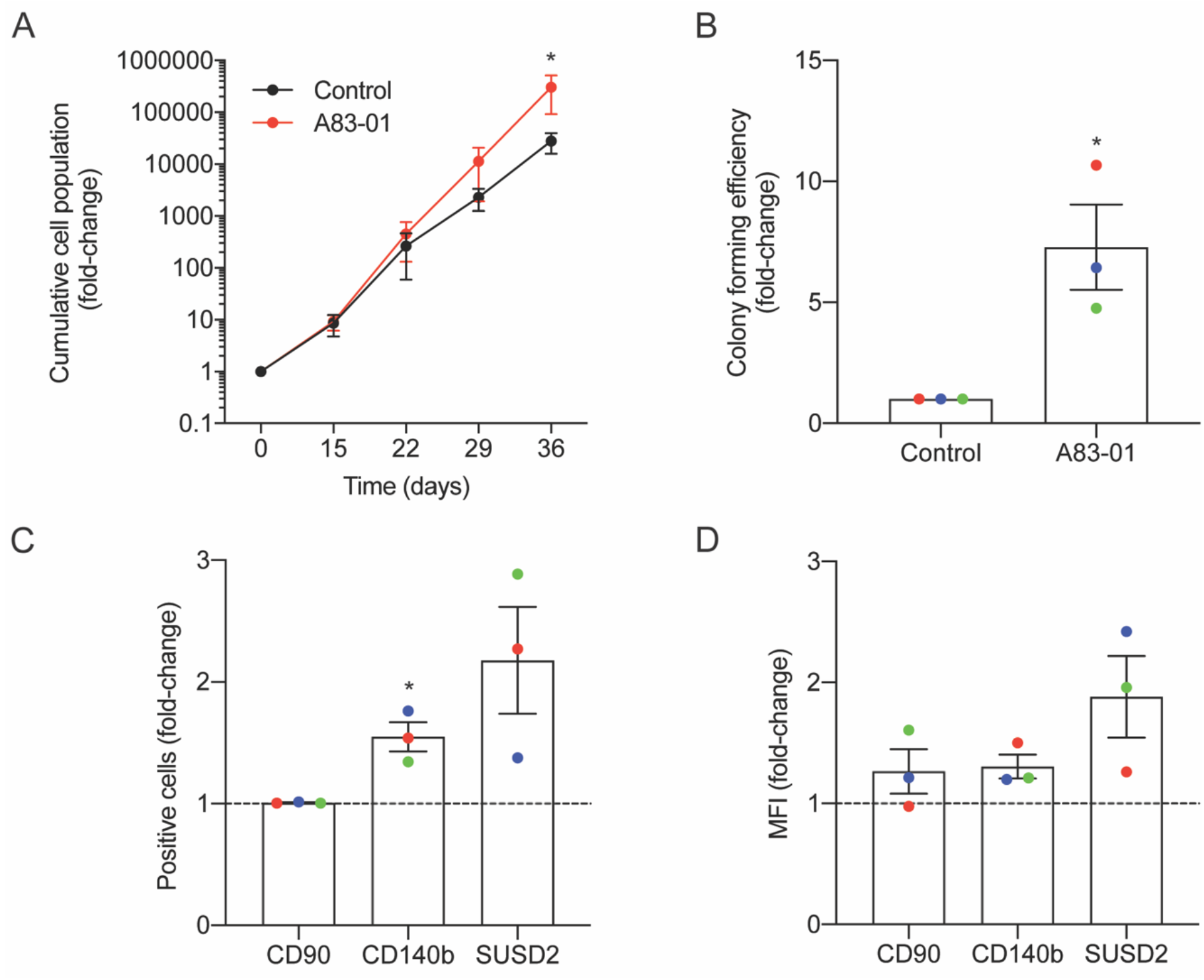
Sustained TGFβ-R inhibition modulates eMSC function and phenotype in culture. Primary human eMSC, isolated using SUSD2 magnetic bead sorting, were cultured with or without 1 μM A83-01 in 5% O_2_ for 36 days. (A) Cumulative cell population in eMSC cultures. Data are mean ± SEM; \**p* < 0.05. Note the logarithmic scale of the Y-axis. (B) Clonogenicity of 3 independent MSC cultures following 29 days culture in SFM with and without A83-01. Surface phenotype assessed by flow cytometry of 3 independent eMSC cultures after 36 days with or without A83-01 shown as (C) % positive cells and (D) MFI for CD90, CD140b and SUSD2. Data are mean ± SEM of fold change over control (shown as dotted line). Individual samples are shown in as different colored dot points. *P < 0.05 from control. Note the % CD90+ cells was ~100% for controls and A83-01-treated eMSC.

### Transcriptional profiling of sustained A83-01-treated eMSC

To explore how cultured eMSC are maintained in a more naïve state upon sustained TGFβ-R inhibition, 3 independent cultures treated with or without A83-01 for 36 days were subjected to RNA-seq. Principal component analysis revealed that the greatest variation in gene expression is accounted for by intrinsic differences between primary cultures. The effect of A83-01 treatment was apparent in principal component 2 (PC2), which accounted for 28% of the variance in gene expression (Fig. 2A). Following Benjamini-Hochberg correction for multiple testing, 1,463 genes were differentially expressed upon A83-01 treatment (Fig. 2B), 759 (52%) of which were up-regulated and 704 (48%) down-regulated. Gene ontology (GO) analysis using DAVID revealed that genes induced upon A83-01 treatment are enriched in 61 biological processes, including ‘regulation of cell growth’ (*p*=7.8×10^−4^) and ‘intracellular receptor signaling pathways’ (*p*=4.5×10^−4^) (Fig. 2C, left panel). Conversely, analysis of downregulated genes yielded GO terms such as ‘cell fate commitment’ (*p*=4.9×10^−4^), ‘collagen catabolism’ (*p*=1.7×10^−13^), and ‘collagen fibril organization’ (*p*=4.6×10^−6^) (Fig. 2C, right panel). Notably, 20 out of the 43 most significantly down-regulated genes upon A83-01 treatment encode extracellular matrix (ECM) components, including various collagen subunits (e.g. *COL1A1*, *COL1A2*, *COL4A1*, *COL4A2*, *COL5A1*, *COL5A2*, *COL6A3*, and *COL8A1*), secreted protein acidic and cysteine rich (*SPARC*), and fibronectin (*FN*) (Fig. S1). Many of the ECM genes repressed by A83-01 are very highly expressed in untreated cells with levels ranging from 342 to 14,586 TPM (Fig. S1). These data suggest that the effect of TGFβ-R blockade on eMSC is mediated, at least in part, by limiting ECM synthesis and deposition in prolonged culture. As shown in Figure S2, several angiogenic, anti-inflammatory, immunomodulatory, antifibrotic and anti-apoptotic genes were significantly upregulated in A83-01-treated cells, in keeping with our previous report.^43^ Further annotation of differentially expressed genes using the KEGG Pathway database underscored the functional differences between A83-01-treated and untreated cultures (Fig. 2D). Notable pathways enriched in eMSC in response sustained TGFβ-R inhibition included the ‘cyclic guanosine monophosphate (cGMP)-protein kinase G (PKG) signaling pathway’, which is implicated in nitric oxide-mediated cardioprotection during acute ischemic preconditioning,^44^, cell growth, and inhibition of apoptosis,^45^ and ‘cytokine-cytokine receptor interaction’, in keeping with the innate paracrine function of MSC.^17, 46^ Conversely, A83-01 repressed genes were enriched in ‘pathways in cancer’, ‘focal adhesion’, and ‘ECM-receptor interaction’. Notably, the ‘PI3K-Akt signaling pathway’ was common to both up- and down-regulated genes. This multifaceted pathway controls key cellular processes by phosphorylating substrates involved in apoptosis, protein synthesis, metabolism, and cell cycle.

**Figure 2.**
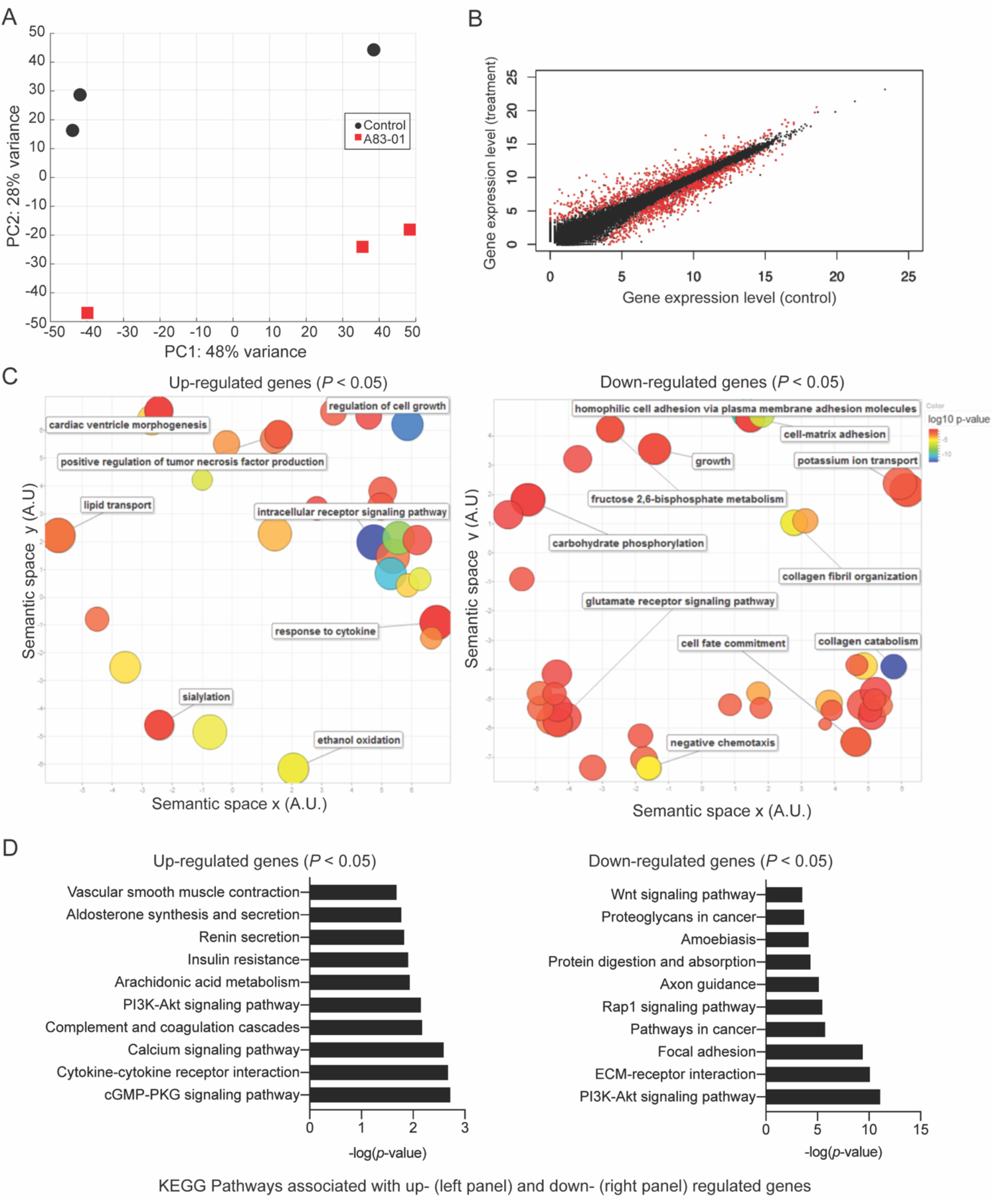
Transcriptomic profile of eMSC upon sustained A83-01 treatment. (A) Principal component analysis of RNA-seq data from three independent primary eMSC cultures treated with or without TGFβ-R inhibitor for 36 days. (B) Average gene expression levels expressed as log2 transformed counts normalized to library size between control and A83-01 treated libraries. Red dots represent significantly differentially expressed genes (*q*<0.05), black dots indicate non-differentially expressed genes. (C) Semantic clustering of significantly overrepresented GO terms (*p*<0.05) of differentially induced and repressed genes (left and right panel, respectively) upon A83-01 treatment. The colour key is shown on the right. The most highly enriched GO categories are indicated in blue. The size of the circles reflects the frequency of the GO term. (D) Enrichment analysis of KEGG pathways associated with up- and down-regulated genes (left and right panels, respectively).

### Chromatin changes induced by sustained A83-01 treatment of eMSC

Dynamic changes in chromatin structure and epigenetic code drive gene expression and ultimately define cell identity.^29^ To map the global changes in the genomic architecture of cultured eMSC in response to sustained TGFβ-R inhibition, 3 independent eMSC cultures treated with or without A83-01 for 36 days were subjected to ATAC-seq, which profiles chromatin accessible regions as a sequencing depth readout.^39^ Based on *q* ≤ 0.05, DESeq identified 5,967 differential ATAC-seq peaks upon A83-01 treatment, 60% of which involved opening of genomic regions and 40% closing of specific loci. Out of 5,967 peaks, 31% and 29% of the opening and closing ATAC-seq peaks, respectively, fell within −10 to +1 kilobases (kb) around transcriptional start sites (TSSs). *RARB* (coding retinoic acid receptor beta, RARβ), *TGFBR3* (transforming growth factor beta receptor 3) and *SUSD2* exemplify genes that showed increased chromatin accessibility at and upstream of their proximal promoters upon A83-01 treatment (Fig. 3A). Cross-referencing with RNA-seq data showed a significant increase in *RARB*, *TGFBR3* and *SUSD2* transcript levels in response to A83-01 treatment (*q*=1.1×10^−33^, *q*=1.2×10^−22^, and *q*=2.7×10^−5^, respectively). Conversely, *CADM1* (cell adhesion molecule 1), *COL1A1* (collagen type I alpha 1 chain), and *WNT5A* are examples of genes repressed in response to A83-01 treatment (*q*=1.9×10^−46^, *q*=1.4 ×10^−9^, and *q*=5.0×10^−30^, respectively). As shown in Figure 3B, downregulation of *CADM1* and *COL1A1* is associated with closure of their proximal promoters whereas silencing of *WNT5A* coincides with closure of a distal enhancer.

**Figure 3.**
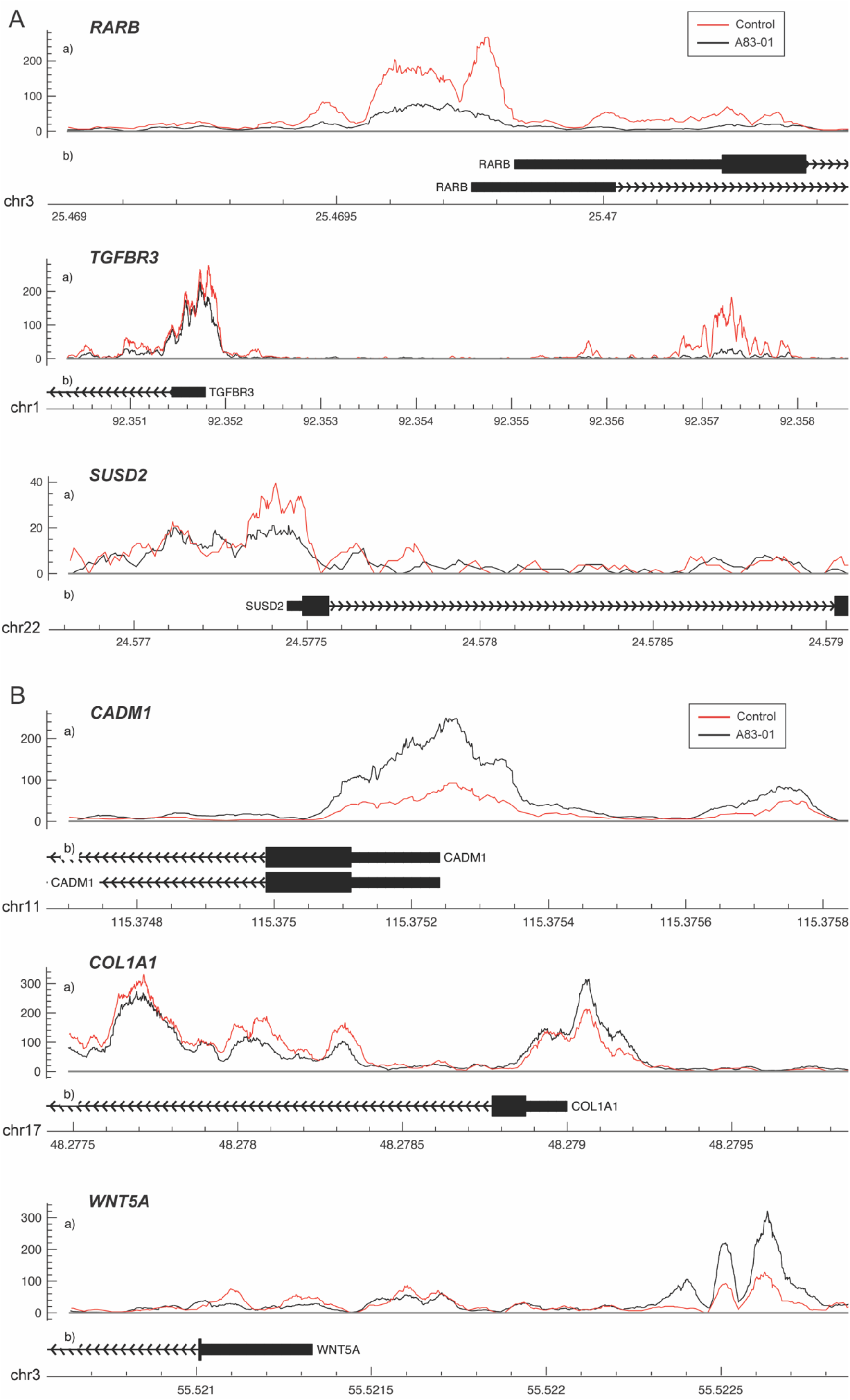
Changes in chromatin accessibility in A83-01 treated eMSC. (A) Representative ATAC-seq peaks showing transition from closing to opening chromatin upstream the promoter of *RARB*, *TGFBR3* and *SUSD2* in response to A83-01. (B) Representative ATAC-seq peaks showing transition from opening to closing chromatin of the proximal promoter of *CADM1* and *COL1A*, and of a distal enhancer of *WNT5A* in response to A83-01. Black and red traces represent untreated and A83-01-treated eMSC cultures. The X-axis shows the genomic location of the ATAC-seq peaks and genes. The Y-axis shows the frequency of Tn5 cutting.

The gain or loss of ATAC-seq peaks upon A83-01 treatment indicates that altered transcription factor (TF) binding drives differential gene expression. However, both activating and repressive TFs can potentially bind at different regulatory sites, rendering it challenging to confidently predict gene expression from dynamic chromatin changes at specific loci alone.^39^ Nevertheless, analysis of 200 genes associated with the most induced or repressed ATAC-seq peaks (within 10 kb of TSSs) revealed a strong association with increased or decreased expression, respectively, upon A83-01 treatment (*p*=1.0×10^−6^) (Fig. 4). Next, we interrogated the ATAC-seq data to gain insight into the *cis*-regulatory landscape that underpins the transcriptional responses of eMSC to A83-01 treatment. *De novo* binding motif enrichment analysis was performed using HOMER on 3,555 opening and 2,412 closing ATAC-seq peaks. This annotation yielded 19 significantly overrepresented motifs in opening peaks (Fig. S3), and 17 overrepresented motifs in closing peaks (Fig. S4). Next, we matched the motifs against canonical TFs that are differentially expressed in cultured eMSC treated with and without A83-01 (Fig. 5A). CCAAT/enhancer binding protein beta and delta (*CEBPB/CEBPD*), *RARB*, RAR-related orphan receptor alpha (*RORA*), and nuclear receptor subfamily 4 group A member 1 (*NR4A1*, also known as *NUR77*) were amongst the most plausible differentially expressed genes of TFs that can bind the enriched motifs in opening ATAC-seq peaks with high affinity (Fig. 5A, B). Conversely, reduced expression of transcription factor 21 (*TCF21*), TGFB induced factor homeobox 2 (*TGIF2*), and nuclear transcription factor Y subunit alpha (*NFYA*) paralleled the loss of their corresponding binding sites in closing loci (Fig. 5A, B).

**Figure 4.**
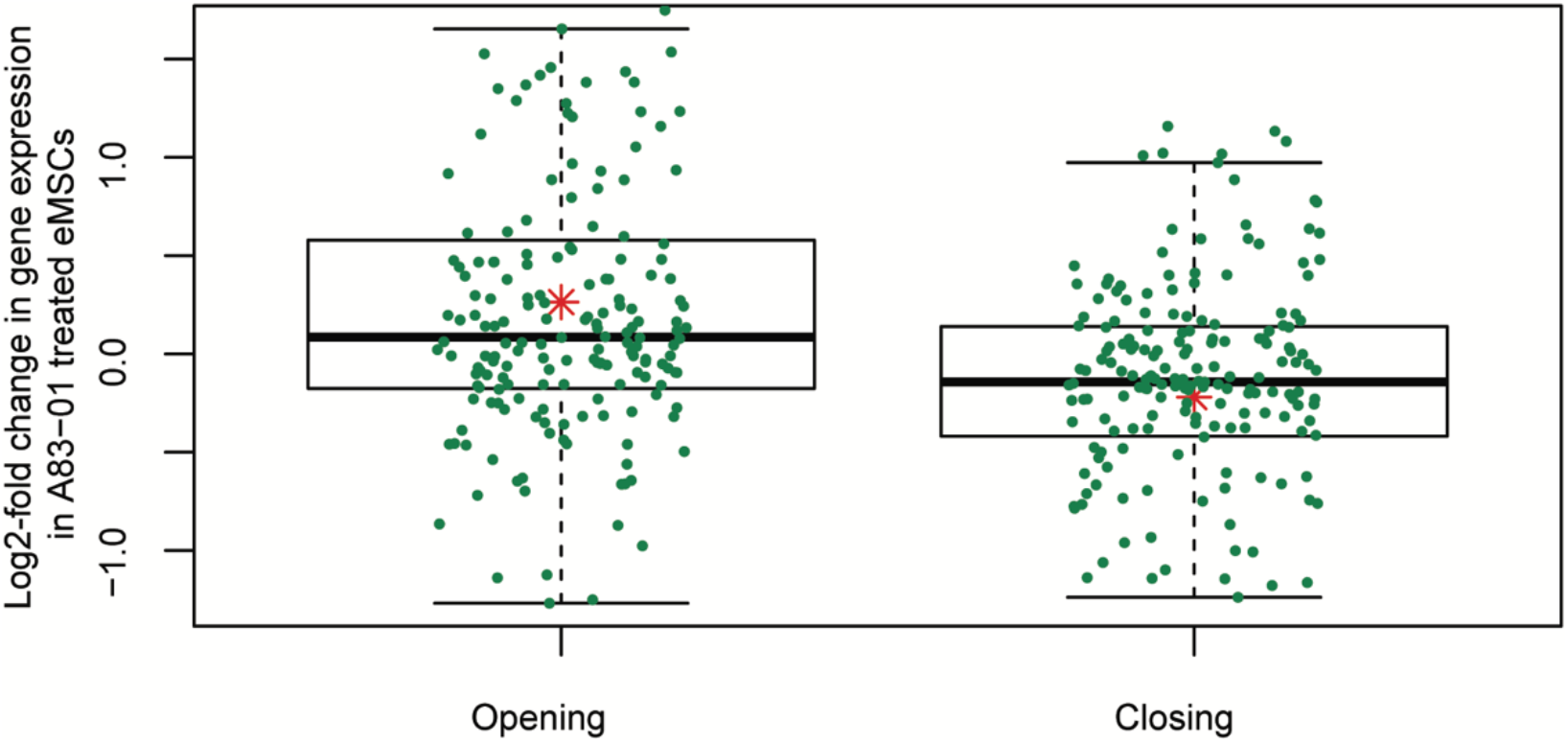
Differential chromatin opening correlates with gene expression changes in A83-01-treated eMSC. Changes in chromatin landscape of eMSC in response to TGFβ-R inhibition correlate with differential regulation of gene expression. Box plots showing increase or decrease in transcript levels of 200 genes (within 10 kb of the TSS) associated with the most open and closed ATAC-seq peaks. Y-axis shows relative changes in transcript levels, expressed as log2-fold change: +*ve* and −*ve* values relate to up- and down-regulated genes, respectively. X-axis shows ATAC-seq peaks clustered in opening and closing peaks. Green dots represent the genes and the red asterisk represents mean log2-fold change (*p*=1.0 × 10^−6^, *t*-test).

**Figure 5.**
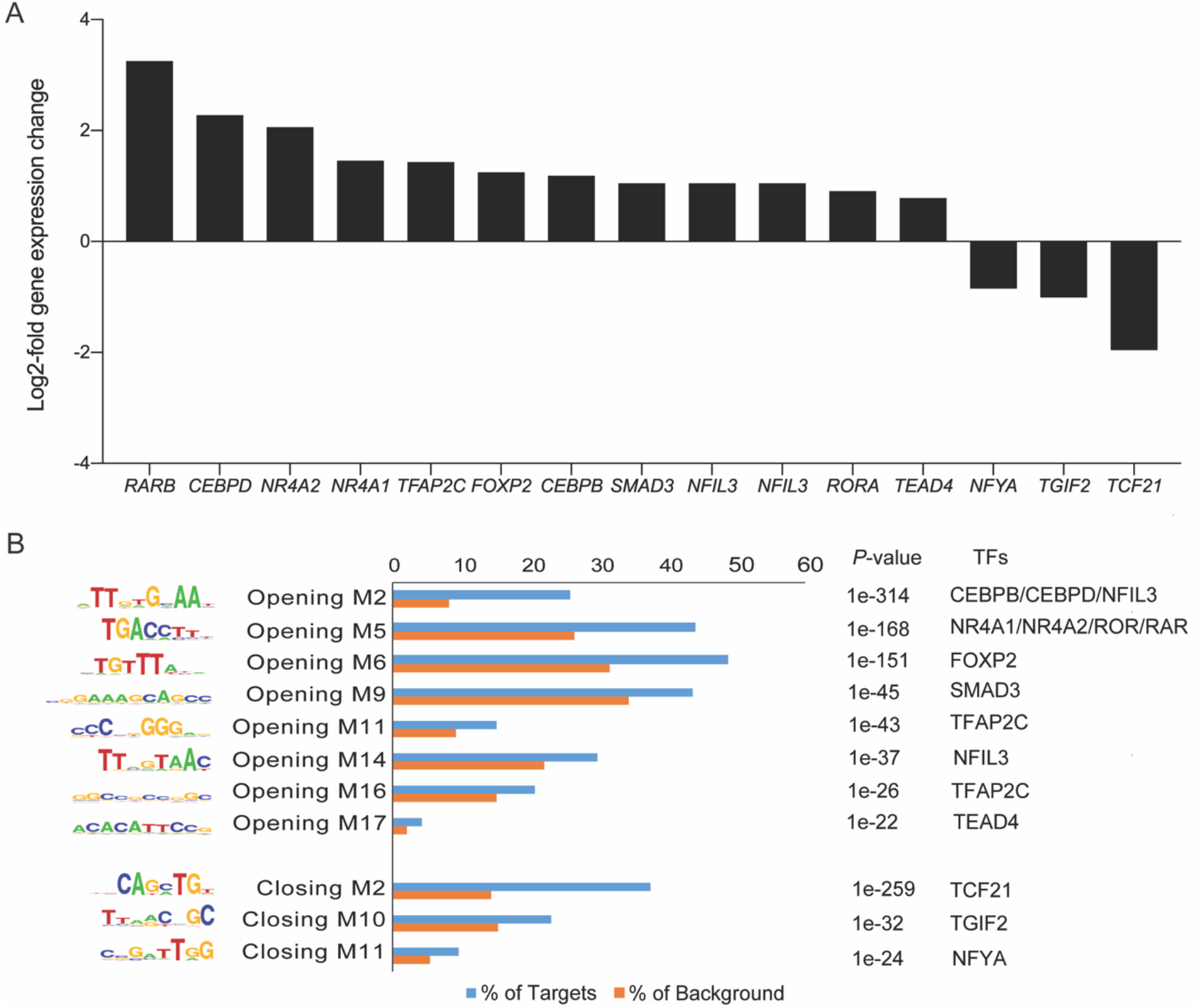
Motif discovery analysis in dynamic genomic regions. (A) Inhibition of TGFβ-R signalling pathway alters expression of genes encoding TFs. Graph shows expression of selected significantly up- and down-regulated TFs (log2-fold change ≥1 and ≤−1). (B) Differentially regulated TFs matched to enriched and depleted short sequence binding motifs. Bar graph showing enriched and depleted binding motifs coupled with the most plausible differentially expressed TFs, based on motif specificity. In the bar graph, the frequency (%) of peaks (blue bars) containing the motif is shown relative to genomic regions randomly selected from the genome (orange bars) (±50 kb from TSS, matching size, and GC/CpG content). *P* indicates the *p*-value of the short sequence binding motifs.

### RARβ is one mediator of A83-01 responses in eMSC

A notable observation was that A83-01 treatment markedly upregulated *RARB* expression (*q*=1.1×10^−33^) (Fig. 5A) in parallel with genome-wide enrichment of RAR/RORA binding sites (*p*=1.0×10^−168^) (Fig. 5B). RARβ is a member of the thyroid-steroid hormone receptor superfamily of nuclear transcription regulators. It binds retinoic acid (RA), the biologically active form of vitamin A.^47^ Further, A83-01 induces multiple other genes implicated in RA signaling (Fig. 6A), including the cellular retinoic acid binding protein 2 (*CRABP2*), as well as known RA target genes (Fig. 6B). *GDF7* (coding growth differentiation factor 7) and *MTSS1* (MTSS I-BAR domain containing 1) exemplify RA target genes that exhibited increased chromatin accessibility and expression in A83-01-treated cells (Fig. 6C). To explore the effect of RARB on A83-01-treated eMSC further, we utilized LE135, a selective RARβ antagonist.^48^ LE135 had no effect on eMSC proliferation in standard cultures not treated with A83-01. However, addition of LE135 to A83-01 treated cultures indicated that RARβ inhibition partially reverses the proliferation advantage conferred by TGFβ-R inhibition (Fig. 6D). In addition, LE135 reduced the clonogenicity of A83-01 treated eMSC (Fig. 6E).

**Figure 6.**
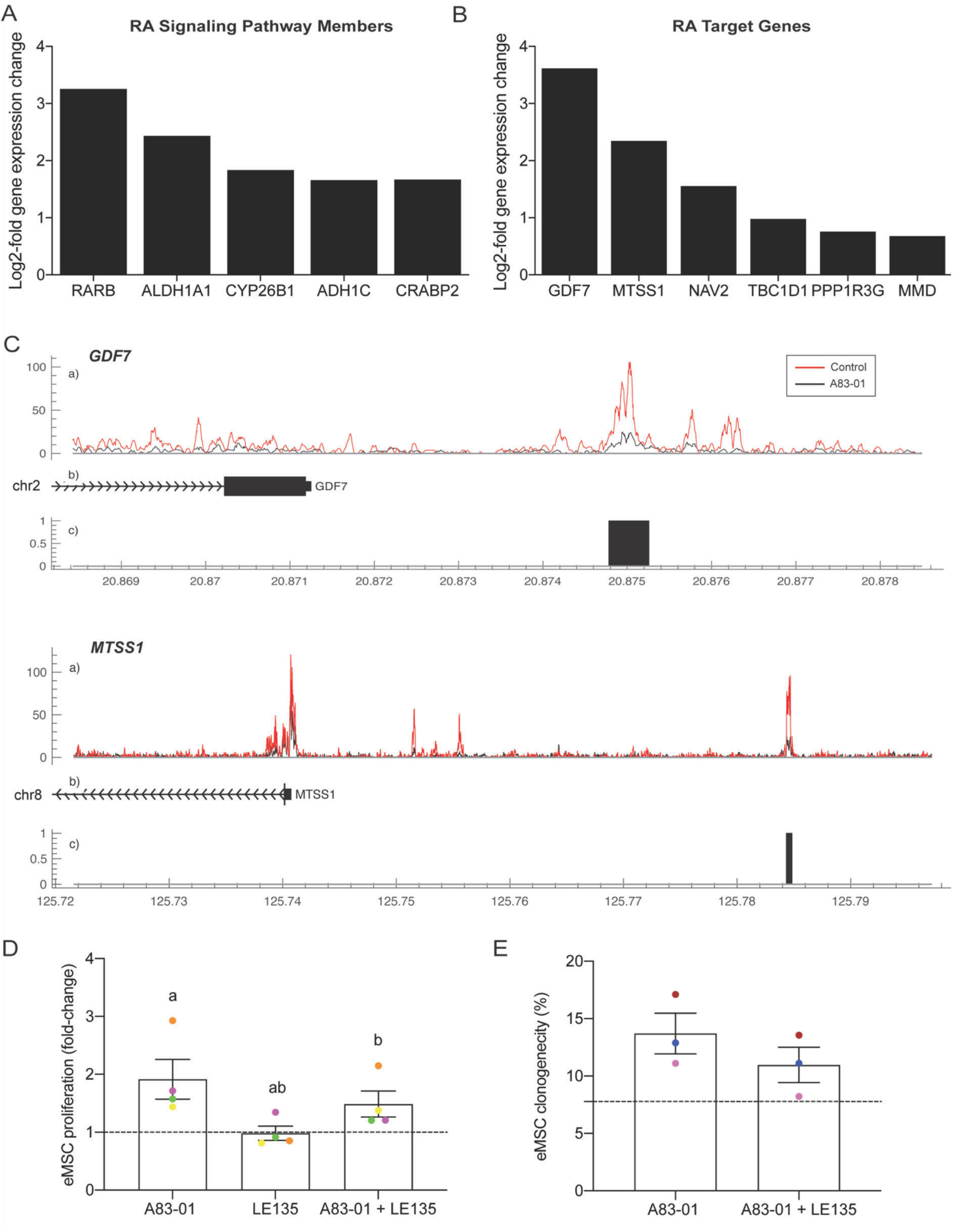
RARβ is one mediator of A83-01 responses in eMSC. (A) A83-01 treatment induces genes implicated in RA signalling and (B) RA downstream target genes. (C) Examples of RA target genes with increased chromatin accessibility. (D) Inhibition of RARβ using a selective antagonist, LE135, partially reverses the proliferation advantage gained upon A83-01 treatment and (E) reduces colony forming efficiency.

## Discussion

This study demonstrates that eMSC cultured under continuous TGFβ-R inhibition are protected against loss of proliferation and clonogenicity upon long-term expansion. The impact of A83-01 on the phenotype of cultured eMSC was underpinned by an altered chromatin landscape and gene expression. More than 1,400 genes were differentially regulated by culturing eMSC in A83-01-containing SFM for 5 weeks, characterized by upregulation of angiogenic, anti-inflammatory, immunomodulatory, antifibrotic and antiapoptotic genes and marked downregulation of ECM genes. The overall signature indicated that A83-01 maintains the expression of genes involved in eMSC paracrine activity while simultaneously inhibiting activation of differentiation genes. Over 3,500 regions of chromatin opened while approximately 2,400 genomic loci closed in response to continuous TGFβ-R inhibition. Motif analysis identified marked enrichment of several putative TF binding sites associated with the more undifferentiated A83-01 treated eMSC, most prominently RAR/ROR. *RARB* expression was also markedly upregulated in parallel with multiple genes encoding signal intermediates in the RA pathway as well as known RA target genes. Functional assays with a selective RARB inhibitor demonstrated an integral role for RA signaling in effecting eMSC responses to prolonged A83-01 treatment. Thus, sustained inhibition of TGFβ-R signaling during eMSC expansion in extended serum-free cultures under physiological O_2_ conditions produces a relatively homogeneous, undifferentiated population of cells with favorable properties for clinical translation.

We reported previously that short-term treatment with A83-01 in SFM (7 days) following extensive culture of eMSC in serum medium (6 passages) increases proliferation and enhances clonogenicity.^23^ Here we report that sustained TGFβ-R inhibition from culture initiation similarly promotes eMSC clonogenicity and proliferation with significant differences observed by passage 3. Prolonged culture under these conditions also increased the percentage of SUSD2- and CD140b-expressing cells and the density of these surface markers on the A83-01-treated cells, but not the representative ISCT marker, CD90, indicating that the eMSC phenotype was retained and spontaneous differentiation attenuated. Our data for these parameters also highlighted intrinsic differences in A83-01 responsiveness between eMSC cultures established from different donors, a well-known feature of primary MSC cultures.^2^ Recently it was shown that body mass index of patients, but not age, correlate inversely with the abundance of clonal SUSD2^+^ eMSC.^49^ These observations raise the possibility that clinical variables, such as obesity, impact on the *in vitro* expandability of eMSC.

The divergent expression of a large number of genes in A83-01-treated cultures suggests either profound transcriptional reprogramming or – perhaps more likely – attenuation of differentiation cues imposed by *in vitro* culture conditions. For example, a striking but not entirely unexpected observation is that A83-01 represses multiple highly expressed genes involved in ECM deposition and collagen metabolism, including 8 collagen subunit genes, *FN1*, *TGFB1* and *SPARC*. During normal wound healing, TGFβ signaling is transiently increased to activate fibroblasts.^50^ Failure to terminate TGFβ signaling following tissue repair results in chronic activation of fibroblasts, massive accumulation of ECM, and fibrosis. Thus, a likely mechanism of A83-01 actions involves prevention of illicit fibroblast differentiation and activation in extended eMSC cultures. In keeping with previous observations,^43^ genes involved in angiogenesis (e.g. *SLIT2*, *HPSE* and *SFRP1*),^51–53^ cell survival (e.g. *ENPP2*, *FAIM2*),^54, 55^ and immunomodulation (e.g. *TLR3* and *IL1R1*)^56, 57^ were also upregulated upon prolonged A83-01 treatment. This gene profile infers that sustained TGFβ-R inhibition leads to expansion of eMSC that are more efficient in reducing inflammation, promoting wound healing and tissue growth, and minimizing fibrosis when transplanted *in vivo*. These functional properties are considered cardinal features of MSC, which were recently renamed as Medicinal Signaling Cells to reflect their paracrine activity.^58^

ATAC-seq analysis revealed that TGFβ-R inhibition in cultured eMSC modifies chromatin accessibility at almost 6,000 genomic regions. The overall pattern, characterized by more opening than closing loci, is in keeping with the less differentiated state of A83-01 treated cells.^29^ A strong correlation was observed between changes in gene expression and differentially chromatin accessibility of promoter regions. For example, opening of chromatin at, and upstream of, the *SUSD2* promoter corresponded to increased abundance of SUSD2^+^ eMSC in A83-01 cultures. Interestingly, SUSD2 has been shown to prevent senescence and cell death in tumor cells,^59^ suggesting its induction is important for expansion of cultured eMSC. Another potential mechanism that promotes survival of A83-01-treated eMSC involves closure of the proximal promoter of *CADM1*, which encodes a potent inhibitor of cell proliferation and migration.^60^ *CADM1* repression is mediated by TWIST1,^61^ a TF upregulated in eMSC in response to A83-01 treatment.^25^ The altered *cis*-regulatory chromatin landscape in response to A83-01-induced TGFβ-R blockade also suggested that silencing of specific TFs may be essential to maintain cultured eMSC in an undifferentiated state. A case in point is the loss of TCF21 binding sites in parallel with marked repression of *TCF21* expression. TCF21 is a member of basic helix-loop-helix family of transcription factors, which orchestrates cell-fate specification, commitment and differentiation in multiple cell lineages during development.^62^ Furthermore, TCF21 has recently been shown to drive fibrosis associated with ovarian and deep infiltrating endometriosis.^63^

Our combined RNA- and ATAC-seq analysis identified the RA pathway as a putative pharmacological target to modulate eMSC in culture. A83-01 markedly upregulated *RARB* expression in parallel with genome-wide enrichment of putative RORA/RAR binding sites. RA plays an important role during embryonic and fetal development and fine-tunes cellular immune response.^64^ In human endometrium, RA signaling is silenced upon differentiation of endometrial fibroblasts into specialized decidual cells ^39^ Induction of RA target genes in A83-01-treated eMSC confirmed that this signaling pathway is activated endogenously in response to sustained TGFβ-R inhibition. Further, the selective RARβ antagonist, LE135, attenuated the cellular responses to A83-01, indicating that increased or sustained RA-RARβ signaling protects eMSC against loss of proliferative capacity and clonogenicity in prolonged culture. Another putative pharmacological target is NR4A1, an orphan nuclear receptor highly induced in A83-01 treated cells. This orphan nuclear receptor exerts pleiotropic regulatory effects on glucose and lipid metabolism,^65^ inflammatory responses,^66^ and vascular homeostasis.^67^ Interestingly, NR4A1 inhibits TGFβ signaling in the nucleus by promoting the assembly of a repressor complex that binds to the promoters of TGFβ target genes.^68^ Because of its role as an endogenous TGFβ inhibitor, small-molecule NR4A1 agonists are under development for the treatment of fibrotic disorders.^68^

In summary, by integrating genome-wide expression and DNA accessibility profiling techniques, this study has advanced our understanding of the *cis*-regulatory DNA landscape and gene networks that safeguards eMSC against spontaneous differentiation and loss of function in prolonged cultures. Analyses of these two large data sets revealed novel pharmacological targets that could be exploited to accelerate clinical translation of autologous eMSC therapies for a variety of reproductive disorders. Furthermore, the data sets constitute a robust resource to interrogate fundamental molecular questions pertaining to human endometrial biology.

## Acknowledgements

We thank the Monash Health Translation Precinct (MHTP) Medical Genomic Facility for the preparation and quality control analysis of RNA and DNA libraries and sequencing.

## Data Availability Statement

RNAseq and ATACseq data from this study have been deposited in the GEO (NCBI) under the accession number GSE146067. The data that support these findings are available from the corresponding author upon reasonable request.

## Graphical Abstract

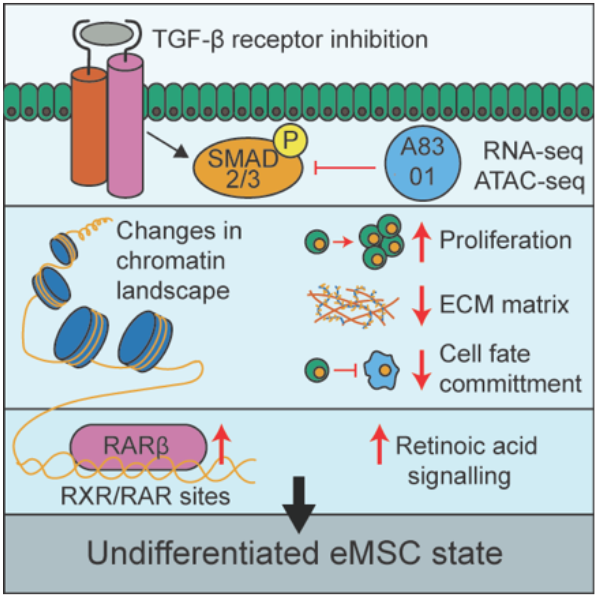

## Supplementary Figure Legends

**Figure S1.**
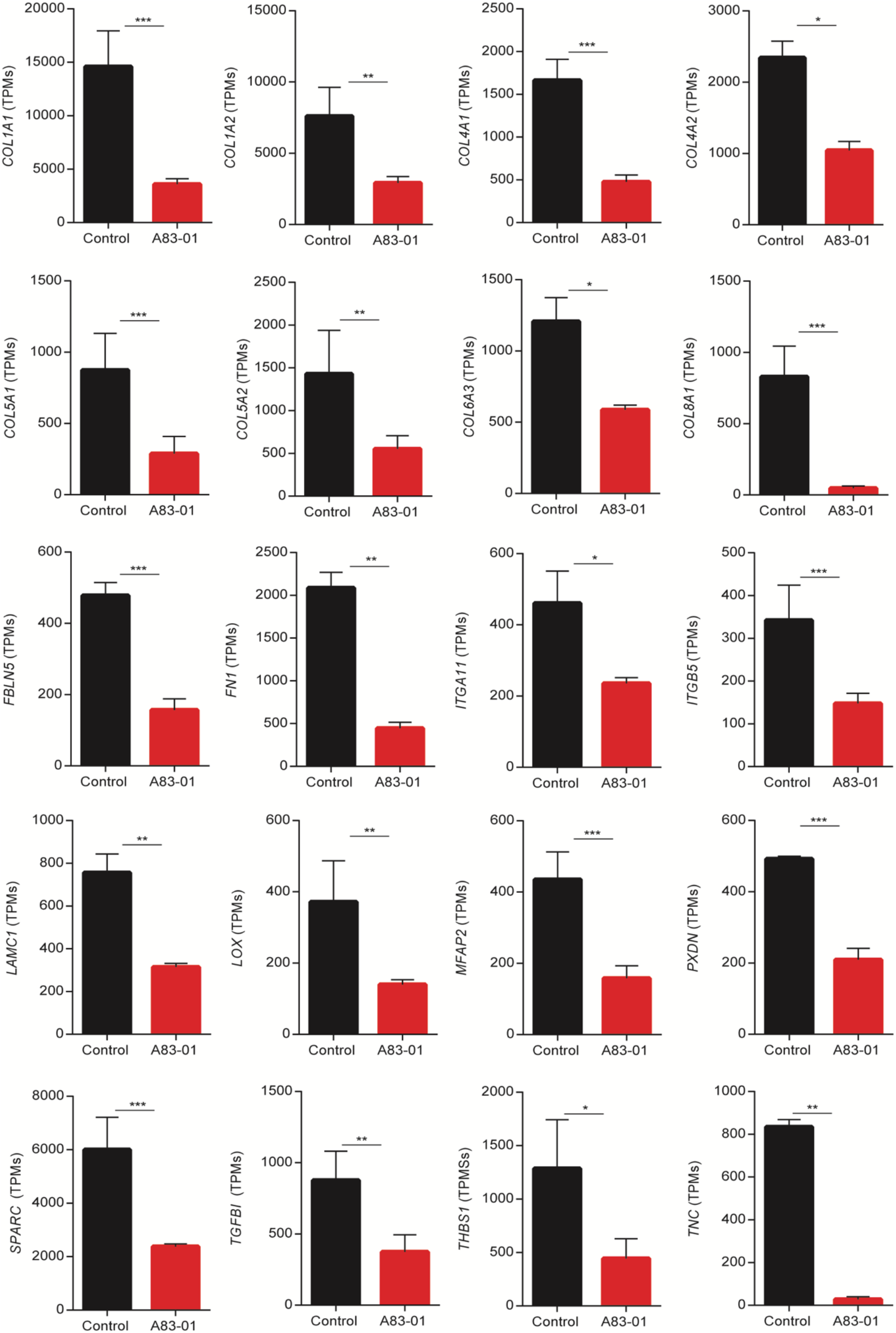
ECM genes down-regulated in response to A83-01 treatment. Graphs showings changes in level of gene expression of the top 16 most abundant ECM genes negatively regulated by TGFβ-R inhibition, represented as changes in TPMs. Data represent mean ± SEM; Y-axis shows TPMs; *indicates *q* < 0.05, ** *q* < 0.01 and *** *q* < 0.001.

**Figure S2.**
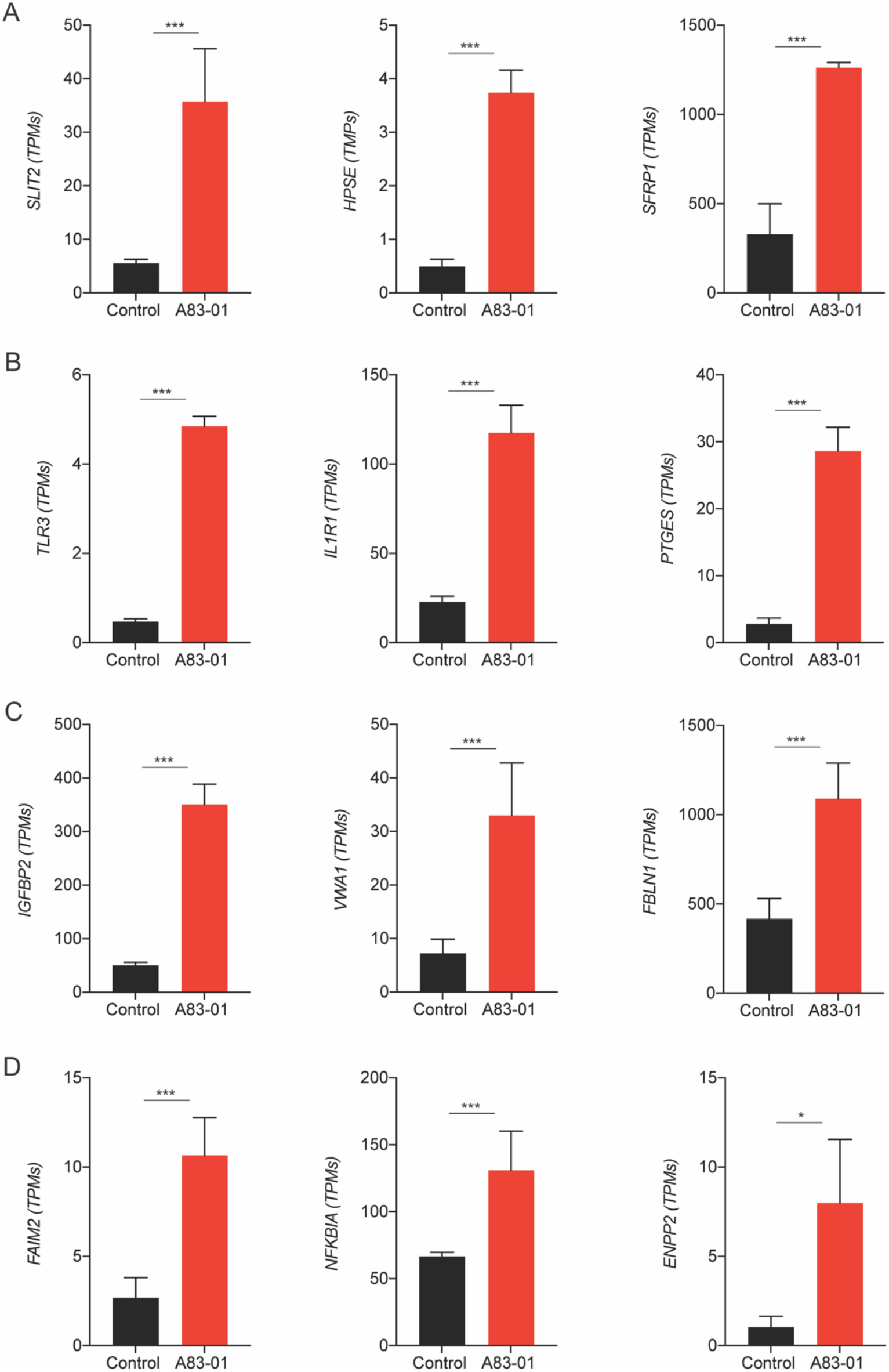
Examples of angiogenic (A) anti-inflammatory, (B) immunomodulatory (C) antifibrotic and (D) antiapoptotic genes up-regulated in response to A83-01 treatment. Graphs show changes in level of gene expression of genes (TPMs) positively regulated by TGFβ-R inhibition. Data represent mean ± SEM; Y-axis shows TPMs; * indicates *q* < 0.05, ** *q* < 0.01 and *** *q* < 0.001.

**Figure S3.**
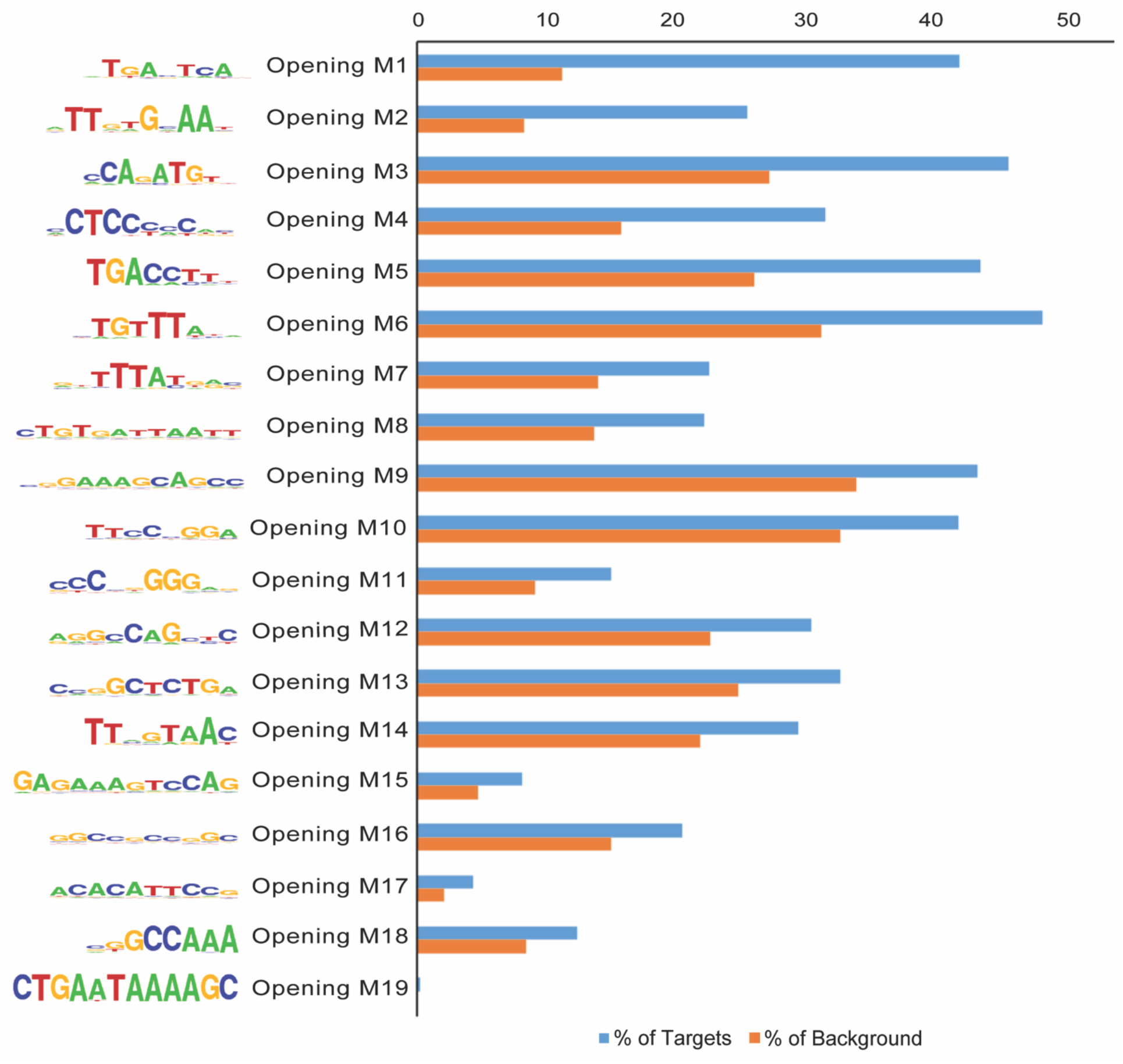
Enrichment of TF binding motifs in opening genomic regions. Total of 19 TF binding motifs enriched in the opening ATAC-seq peaks. The frequency (%) of peaks (blue bars) containing a given motif is shown relative to genomic regions randomly selected from the genome (orange bars) (±50 kb from TSS, matching size, and GC/CpG content.

**Figure S4.**
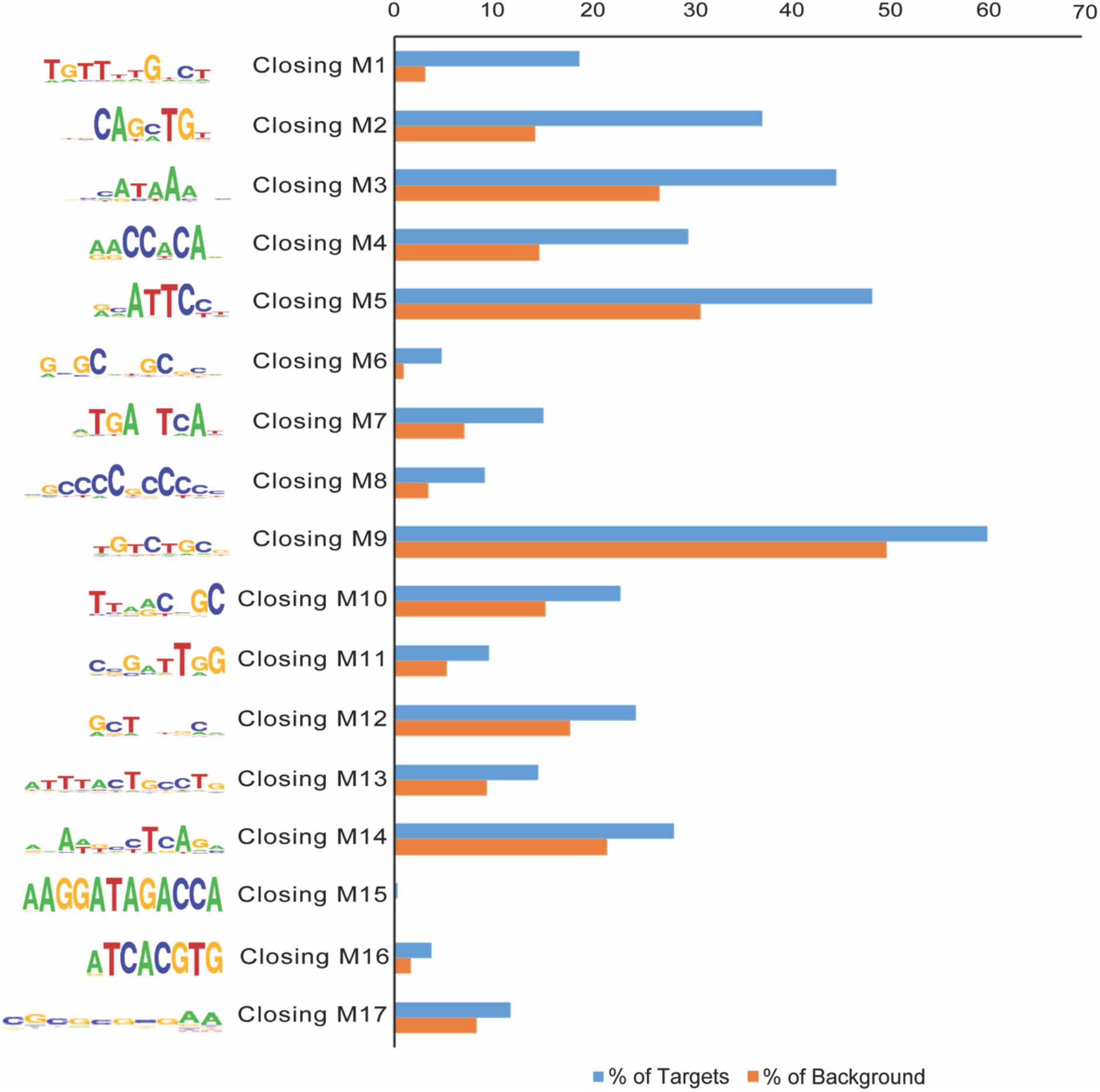
Depletion of TF binding motifs in closing genomic regions. Total of 17 TF binding motifs enriched in the closing ATAC-seq peaks. The frequency (%) of peaks (blue bars) containing a given motif is shown relative to genomic regions randomly selected from the genome (orange bars) (±50 Kb from TSS, matching size, and GC/CpG content).

